## Supplemental Figure 1 for "Attenuated influenza virions expressing the SARS-CoV-2 receptor-binding domain induce neutralizing antibodies in mice"

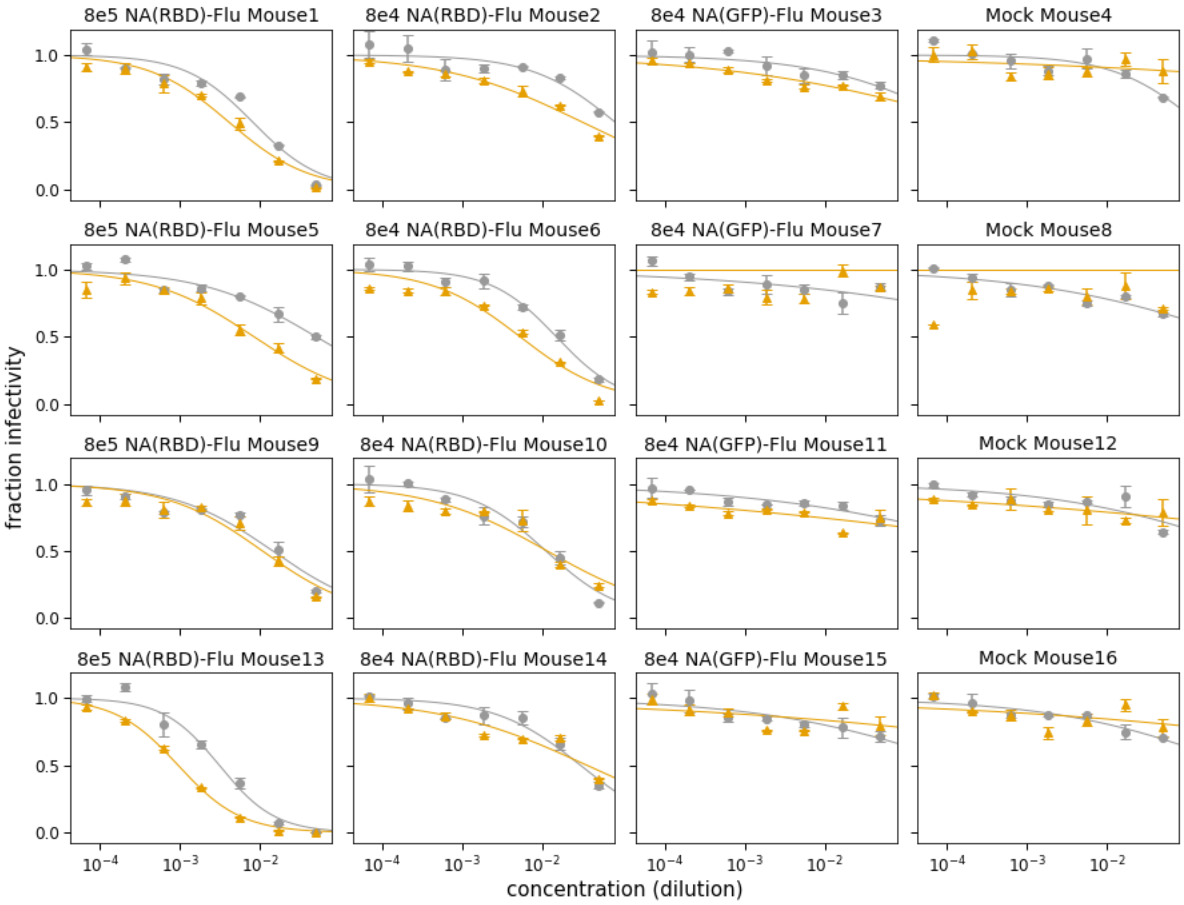


**Supplemental Figure 1.** Neutralizing antibodies are present in mice at 14- and 21-days post inoculation with ΔNA(RBD)-Flu. Neutralization against SARS-CoV-2 pseudotyped-lentivirus is compared for sera collected at 14- (grey) and 21- (orange) days post infection with either 8e5 (RBD)-Flu, 8e4 ΔNA(GFP)-Flu, or mock infected. Each graph represents a single mouse, curves are plotted to compare neutralization at each timepoint. Fraction infectivity was calculated by normalizing the luciferase reading for each sample by the average of the two no-serum control wells in the same row. All neutralization curves represent the mean and standard error of two replicate curves run on the same 96-well plate. Neutralization curves were plotted using the *neutcurve* Python package (https://jbloomlab.github.io/neutcurve/, 0.3.1), which fits a two-parameter Hill curve.
